# Yeast grown in continuous culture systems can detect mutagens with improved sensitivity relative to the Ames test

**DOI:** 10.1101/2020.06.15.152371

**Authors:** Joseph Y Ong, Julia T Pence, David C Molik, Holly V Goodson

**Author notes:** Corresponding author: Holly V Goodson.

## Abstract

Continuous culture systems allow for the controlled growth of microorganisms over a long period of time. Here, we develop a novel test for mutagenicity that involves growing yeast in continuous culture systems exposed to low levels of mutagen for a period of weeks. In contrast, most microorganism-based tests for mutagenicity expose the potential mutagen to the biological reporter at a high concentration of mutagen for a short period of time. Our test improves upon the sensitivity of the well-established Ames test by at least 20-fold for each of two mutagens that act by different mechanisms (the intercalator ethidium bromide and alkylating agent methylmethane sulfonate). To conduct the tests, cultures were grown in small, inexpensive continuous culture systems in media containing (potential) mutagen, and the resulting mutagenicity of the added compound was assessed via two methods: a canavanine-based plate assay and whole genome sequencing. In the canavanine-based plate assay, we were able to detect a clear relationship between the amount of mutagen and the number of canavanine-resistant mutant colonies over a period of one to three weeks of exposure. Whole genome sequencing of yeast grown in continuous culture systems exposed to MMS demonstrated that quantification of mutations is possible by identifying the number of unique variants across each strain but with lower sensitivity than the plate-based assay. In conclusion, we propose yeast grown in continuous culture systems can provide an improved and more sensitive test for mutagenicity.

## Introduction

Continuous culture systems allow for the long-term, controlled growth of microorganisms or cultured cells. Consequently, they have been used for many scientific and industrial uses, including producing biologics like small molecules [1–3] and recombinant proteins [4,5]; assessing the growth rate [6–9] or metabolism [10,11] of microorganisms or cultured cells under defined conditions; and for studying evolution [12–14]. Continuous culture systems typically operate in one of two modes: a chemostat, where a limited amount of nutrients are constantly added to the culture, or a turbidostat, where the culture density is kept constant [15]. Here, we present a test for using yeast grown in chemostats as a novel test for mutagenicity.

*In vitro* tests for mutagenicity typically expose bacteria or cultured mammalian cells to the suspected mutagen and assay for gross DNA damage, such as micronuclei or chromosomal aberrations, or for mutations of particular genes, such as HPRT with Chinese hamster ovary cells (reviewed by [16]). One of the common tests for mutagenicity is the Ames test, a bacteria-based test that uses reporter *Salmonella typhimurium* lines to assess mutagenicity [17,18]. In the original form of this assay, the bacteria have a mutation in histidine synthesis gene *hisG46* and are auxotrophic to histidine. The bacteria are exposed to the potential mutagen and plated in media with limited histidine, and the number of colony forming units (bacteria that mutated and reverted to being prototrophic to histidine) are counted and assessed as a measure of mutagenic potential. Similar assays have been developed for *Escherichia coli* and tryptophan synthesis [18,19].

While inexpensive and easy to perform, the Ames test does have limitations. First, the Ames test exposes the bacteria to high concentrations of a single test compound over a short period of time, usually 48 hours [20], as low concentrations of mutagen may lead to the control and experimental groups having similar numbers of mutant colonies, leading to an ambiguous result [21]. However, likely human exposure to suspected mutagens is expected to be at low concentrations over a very long period of time, such as drinking water contaminated by a mixture of pollutants, each present in very small concentrations [22]. In assessing the safety of such water, it is generally assumed that effects of potential mutagens are both linearly related to concentration and additive, but whether these assumptions are correct is difficult to determine with the present technologies. Moreover, the Ames test and similar assays based on mutation reversion require that the strain be matched to the type of mutagen being detected: a strain with a point mutation is used to detect compounds (e.g., alkylating agents) that act by changing DNA sequence, while a strain with a frameshift is needed to detect intercalators. Consequently, there is a need for an improved test for mutagenicity, one that has improved sensitivity and can assess multiple types of mutagens while remaining inexpensive.

Here, we present yeast grown in continuous culture systems as a novel method of mutagenicity testing. Yeast are an ideal organism for continuous culture systems because they grow robustly at room temperature, are well-established model organisms, perform well in continuous culture systems [12,23], and have been previously functionalized as biosensors [24]. Moreover, as yeast are eukaryotes, their metabolisms and DNA repair pathways are more similar to that of humans, potentially providing an advantage for mutagenicity testing when compared to bacteria. Our approach allows for the assessment of yeast exposed to mutagen over the span of approximately 1-2 weeks, permitting us to assay low concentrations of mutagen over long periods of time. With this approach, we were able to detect levels of the mutagens methyl methanosulfate (MMS; an alkylating agent) and ethidium bromide (EtBr; an intercalator) at concentrations at least an order of magnitude lower than the Ames test via a canavanine plate-based assay. We were also able to detect point mutations in yeast grown in MMS via whole genome sequencing, though with less sensitivity. Altogether, we demonstrate that a yeast-based chemostat approach is a viable method of determining the mutagenicity of compounds at low concentrations.

## Materials and Methods

### Preparation and assembly of chemostats and media containers

Chemostats were constructed by drilling holes into standard 15 mL polypropylene conical tubes (VWR), with one hole at the bottom for the addition of an air inlet and one hole at the 5 mL mark for the addition of a waste outlet. To the cap were drilled two adjacent holes for the addition of media. To the holes in the cap and at the bottom of the culture chamber were attached 1/16” barbed ports, and to the waste outlet was attached a 1/8” barbed port. These ports were sealed in place with commercial two-part epoxy and allowed to set over 48 hours, per manufacturer’s instructions. One of the ports on the cap was sealed with a small piece of 1/16” tubing connected to a sealed plug. Prepared chemostats were placed in a sealed beaker and autoclaved. Autoclaving at 120°C for 15 minutes showed no signs of degradation in the epoxy or elsewhere, even after repeated autoclave cycles.

Two holes were drilled through the caps of 500 mL Pyrex jars and 1/8” female luers were attached through both of them and sealed with epoxy. One female luer was sealed with a male luer plug. Media was prepared and 200 mL was transferred into the glass jar. The modified lid was placed over the jar and a length of 1/16” tubing was fed through the open female luer until the tubing comfortably reached the bottom of the jar. The opening of the female luer was closed with autoclave tape and aluminum foil. The media was then autoclaved at 120°C for 15 minutes.

Peristaltic pumps (SP100 peristaltic pumps; APT Instruments, A/C, 15 rpm) were used to pump media into the chemostat. The inner tubing (silicon, 0.8 mm in diameter) was autoclaved for 120°C for 15 minutes. An aquarium pump (Marina 200 Fish Tank Aquarium Air Pump) was used for aerating the chemostats. Before assembly, 0.2 micron filters (VWR, USA) were attached to the two outlets of the air pump to sterilize the air leaving the aquarium pump and entering the culture chamber. Each filter was connected with sterile 1/8” tubing to a four-way manifold. Four-way manifolds were sterilized with 70% ethanol and allowed to dry immediately before use. Stopcocks on the four-way manifold allowed for the adjustment of the airflow to a gentle rate of about 2 to 3 bubbles per second.

To assemble the culture chambers, the inner tubing of the peristaltic pumps was positioned within the housing of the pump and secured. A length of 1/16” tubing about 30 cm long was used to connect the peristaltic pump to the cap of the culture chamber. Another length of 1/16” tubing was used to connect the bottom of the culture chamber to the four-way manifold. A length of 1/8” tubing was connected to the outlet port and the free end of the tubing was allowed to feed into a sterile 50 mL conical tube sealed with aluminum foil. To the culture was added about 3 mL of an overnight culture of yeast grown from one colony of DBY10148 [25] (*Saccharomyces cerevisiae*, gift of David Botstein) in synthetic complete media (described below) with ampicillin. DBY10148 is a chemostat-adapted strain of yeast that does not clump or stick to the sides of the continuous culture tube. Other strains could potentially be used, but for proper performance in the chemostats, it is imperative to avoid strains that clump and/or stick to the sides of the tubes.

The peristaltic pumps were plugged into a power strip that was controlled by an electronic outlet timer set to turn on the pumps for one minute every hour. After controlling air flow rate through the culture chambers, all the peristaltic pumps were turned on to fill the continuous culture systems to the appropriate volume before allowing the electronic timer to dictate culture feeding times. Similarly, whenever media was changed, the pump corresponding to that chemostat was allowed to run until the media had cleared the empty space in the tubing and reached the culture chamber.

### Preparation of synthetic media

Synthetic complete media was prepared by adding 20 g D-glucose (VWR, USA), 6.7 g Difco(^tm^) Yeast Nitrogen Base w/o Amino Acids (Bioworld, Grinnell, IA), and 2.0 g Drop-Out Mix Complete (US Biological, Swampscott, MA) to 1 L of DI water. We used synthetic complete media instead of standard rich media (e.g., YPD) because clumping problems sometimes appeared when growing cultures in YPD. The solution was divided among five 500 mL jars (200 mL of media each) and capped with a modified lid. One port of the lid was sealed with a screw-cap luer, and through the other was fed a length of 3 mm in diameter tubing so that the tubing comfortably reached the bottom of the jar. The opening through the tubing port was sealed with autoclave tape and aluminum foil. The media was autoclaved at 120°C for 15 minutes and stored at room temperature until use. Ampicillin (Sigma, St. Louis, MO; final concentration 100 mg/L), methyl methanesulfate (MMS; Sigma; used freshly prepared from stock bottle), and ethidium bromide (EtBr; Fischer, Fair Lawn, NJ; aqueous stock solution: 10. mg/mL) were added through the screw-cap luer port before use. Aluminum foil was wrapped around media containers to protect EtBr from degradation from light, and media was stored away from direct sunlight.

### Preparation of canavanine plates

Canavanine plates were made by adding 20 g D-glucose (VWR), 6.7 g Difco(^tm^) Yeast Nitrogen Base without Amino Acids (Bioworld), 2.0 g Drop-Out Mix Synthetic Minus Arginine without Yeast Nitrogen Base (US Biological), and 20 g agar (Research Products International (RPI); Mt. Prospect, IL) to 1 L of DI water and autoclaving at 120 °C for 15 minutes. When the mixture had cooled to about 55°C, canavanine sulfate (Sigma; final concentration: 30 mg/L) and ampicillin (Sigma; final concentration: 100 mg/L) were added and the mixture allowed to stir before pouring into 15 × 100 mm plates (about 25 mL per plate). Plates were allowed to set overnight and stored at 4°C until use.

### Plating of Yeast

To collect a yeast sample, the outlet port was switched to feed into a sterile 15 mL conical tube and left for 2-3 hours to collect about 1-2 mL of yeast culture. Five hundred microliters of the collected yeast culture were saved as a glycerol stock (25% by volume glycerol) and stored at -80ºC. The culture density (OD600) was measured and 1 OD unit of yeast was plated on canavanine plates. Lower amounts of yeast (e.g., 0.5 OD units) were plated as the number of canavanine-resistant colonies increased to maintain evenly spaced and countable colonies, and the final numbers were adjusted to account for this dilution. Sterile PBS was used to bring the volume of yeast up to 200 μL if necessary and the yeast culture was spread on the plate with sterile glass beads. The plates were incubated at 30ºC for two or three overnights, after which canavanine-resistant colonies were manually counted. The source data, including the number of counted canavanine-resistant colonies for all experiments, are available in S1_Table.

### Yeast DNA extraction

For genomic DNA analysis, yeast were streaked out from a glycerol stock onto a YPD agar plate (50 g Difco(^tm^) YPD (BD, USA) and 20 g agar (RPI) dissolved in 1 L of DI water, autoclaved at 120°C for 15 minutes, and poured into 15 × 100 mm plates, about 25 mL per plate) and three separate colonies were selected and grown in 40 mL of YPD (50 g Difco(^tm^) YPD dissolved in 1 L of DI water and autoclaved at 120°C for 15 minutes) in a 50 mL conical tube at 37°C overnight with gentle shaking. After centrifuging the overnight culture at 4,000 rpm for 5 minutes to pellet the yeast, the supernatant was discarded and the yeast were resuspended in 3 mL of 0.9 M sorbitol and 0.1 M Na_2_EDTA (pH 7.5). To this was added 0.1 mL of 2.5 mg/mL solution of Zymolyase 100T (Sigma) in 0.9 M sorbitol, and the mixture was incubated for one hour at 37°C with gentle shaking. The mixture was centrifuged at 4,000 rpm for 5 minutes, and the supernatant was discarded.

The cell pellet was first resuspended in 5 mL of 50 mM Tris-Cl and 20 mM Na_2_EDTA (pH 7.4), then 0.5 mL of 10% (w/v) SDS was added. The mixture was carefully stirred using a glass pipette and incubated at 65°C in a water bath for 30 minutes, with a gentle mixing at 15 minutes. Potassium acetate (1.5 mL of 5 M) was added, mixed thoroughly, and left on ice for one hour, with thorough mixing every 15 minutes to yield a viscous white mixture. The mixture was centrifuged at 10,000 rpm in a Sorvoll SS-34 rotor for 10 minutes to yield a thick, wide pellet.

The supernatant was carefully transferred into a clean Oak Ridge tube and to the supernatant was added two volumes (about 10 mL) of absolute ethanol. This solution was centrifuged at 5,500 rpm for 15 minutes, and the supernatant was carefully decanted. The Oak Ridge tube was inverted to allow the pellet to air-dry for about ten minutes before the pellet was resuspended by gentle mixing in 3 mL of TE buffer (10 mM Tris-Cl, 1 mM Na_2_EDTA, pH 7.4). The solution was transferred into two 1.5 mL Eppendorf tubes and centrifuged at 13,000 rpm at 4°C for 10 minutes to yield a small pale white or pale white-yellow pellet. The supernatant from both tubes was recombined into one 15 mL conical tube, and to this new tube was added 7.5 µL of RNAse A (20 mg/mL from Sigma) and mixed gently. The solution was incubated at 37°C for 30 minutes with gentle shaking.

Two volumes (about 6 mL) of isopropanol were added into the 15 mL conical tube and gently inverted to yield a loose “cobweb” of DNA. The DNA was pelleted by centrifugation at 4,500 rpm at 4°C and the supernatant was carefully decanted. The DNA pellet was resuspended by the addition of 200 µL of DI water, transferred to a 1.5 mL Eppendorf tube, and two volumes (400 µL) of absolute ethanol was added to the tube. After gentle inversion, the tube was centrifuged at 5k rpm for 5 minutes at 4°C, the supernatant was removed by careful pipetting, the tube was left inverted for 15 minutes so the remaining ethanol would air-dry, and the DNA resuspended in 50 µL of DI water. The purity and integrity of the genomic DNA was assessed by Nanodrop and TE-agarose gel electrophoresis, and the DNA was stored at -20°C until sequencing.

### Counting variants via whole genome sequencing

Sequencing of DNA samples as prepared above was performed at the University of Notre Dame Genomics and Bioinformatics Core Facility with an Illumina MiSeq desktop sequencer. Twenty-seven genomes (the starting strain of yeast and two time points with four conditions per time point; all in triplicate) were prepared with an Illumina Nextera DNA Sample Preparation Kit with V3 600 cycle kit to generate 300 base pair paired-end reads. Reads were processed through the University of Notre Dame Center for Research Computing. The CEN.PK2-1 reference genome [26] (GenBank JRIV00000000.1) or S288C reference genome (GenBank GCA_000146045.2) was indexed with BWA [27] (version 0.7.17) and Illumina reads were aligned to the CEN.PK2-1 reference genome to generate SAM files. SAM files were converted to BAM files, and BAM files were sorted and indexed using SAMtools [28] (version 1.9).

For variant calling with VarScan, an MPILEUP file was generated with SAMtools and variants were called with somatic variant calling with VarScan [29] (version 2.4.3) using one colony of the initial culture of yeast as baseline (“normal”) and the yeast grown in continuous culture as a comparison (“tumor”). Single nucleotide polymorphisms (SNPs) were manually filtered to exclude ambiguous calls (that is, SNPs where two different alleles were called at one genomic position) and to exclude SNPs with a p-value greater than 0.05 (as determined by VarScan via Fisher’s Exact Test). For variant calling with Mutect2, a dictionary for the CEN.PK2-1 reference genome was created with Picard tools (https://github.com/broadinstitute/picard, version 2.22.4), and the reference genome was indexed using SAMtools. Read groups were added to sorted BAM files with Picard tools. One of the Day 0 colonies, designated as the baseline (“normal”) was given sample ID 1; all the remaining 26 colonies were given sample ID 20. To call variants, Mutect2 from GATK [30] (version 4.1.7.0) was used in a similar manner as VarScan2 to call variants. SNPs were also manually filtered to exclude ambiguous calls and to exclude SNPs with 2 or fewer reads supporting that call.

## Results

### Construction of chemostat devices

We designed an inexpensive continuous culture system that would allow yeast to grow undisturbed over a long period of time (weeks). For the growth chamber, we used a conventional 15 mL conical tube (**Fig 1A**). A hole was drilled at the bottom of the tube and an inlet was inserted and secured with epoxy. Tubing was used to connect the bottom inlet to an aquarium pump that gently bubbled filtered air into the culture, aerating the culture. This aquarium pump was always on when the cultures were growing. A similar hole was drilled into the cap of the tube, and a peristaltic pump delivered synthetic yeast media (with or without mutagen) into the culture via this inlet. A timer turned this pump on for one minute every hour, allowing approximately 0.67 mL of media into the culture. Finally, a hole was drilled into the side of the conical tube at approximately the 5 mL line and an outlet valve was inserted. When a sample was being collected, this outlet valve was directed into a collection tube; otherwise, the outlet valve led to a waste container. Accounting for bubbling, the yeast culture volume within the chemostat was about 3 mL, and excess culture would flow out of the outlet tube during feeding times, as the volume of the culture remained constant. To start a chemostat, an overnight culture of yeast was diluted down to 0.5 OD600 and split evenly (∼3 mL each) among the culture systems. Four or five of these cultures were run in parallel. For our experiments, we used MATa yeast strain DBY10148, a strain adapted for use in continuous culture systems [25].

**Fig 1.**
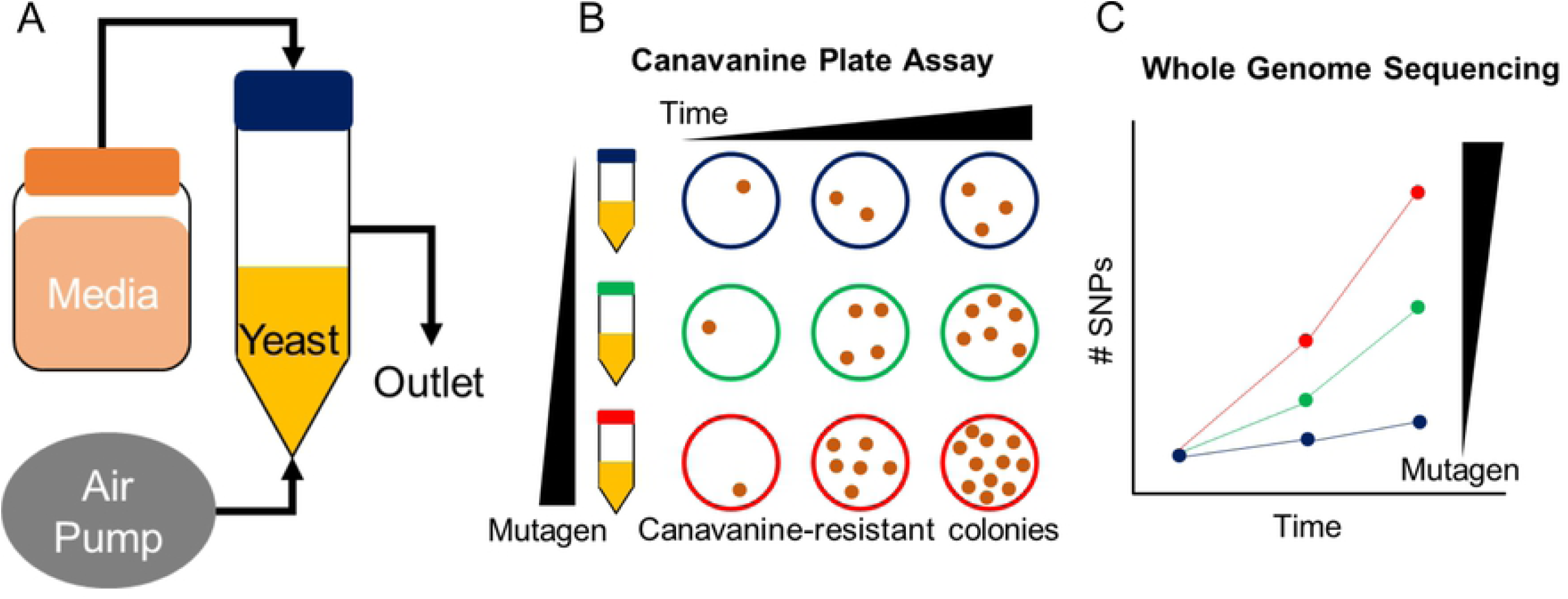
Overview of technique for using yeast as a biosensor for mutagenicity. (A) Schematic of a chemostat culture system demonstrating the positions of inlet port for aeration and for media input at regular intervals and the outlet port for collecting waste and yeast samples. (B) Schematic of the canavanine-based plate assay and (C) whole genome sequencing-based assay used to determine mutagenic potential over time and as a function of mutagen concentration. In (B), the number of cavananine-resistant yeast colonies serves as a measure of mutagenic potential, whereas in (C), the number of SNPs or other mutations serves as a measure of mutagenic potential.

### Use of yeast chemostats to measure mutagenicity of the alkylating agent MMS

To assess the ability of these chemostats to detect lower concentrations of mutagens, we first focused on testing the DNA methylating agent methylmethane sulfonate (MMS) [31]. Given that the limit of detection for the Ames test is approximately 2.2 - 4.4 mg MMS/L [32,33], we began by growing yeast in four parallel chemostats with 0, 0.1, 1.0, and 10 mg MMS/L (**Fig 2A**). About every three days for a month, yeast from each chemostat were plated on synthetic media plates without arginine and supplemented with canavanine. Canavanine is a non-proteogenic amino acid structurally similar to arginine and toxic to wild-type yeast. Yeast with mutations in the *CAN1* gene may fail to import canavanine and consequently survive [34]. As canavanine resistance correlates with mutations in the *CAN1* gene, we counted canavanine-resistant yeast colonies as a measure of mutagenicity (**Fig 1B**).

**Fig 2.**
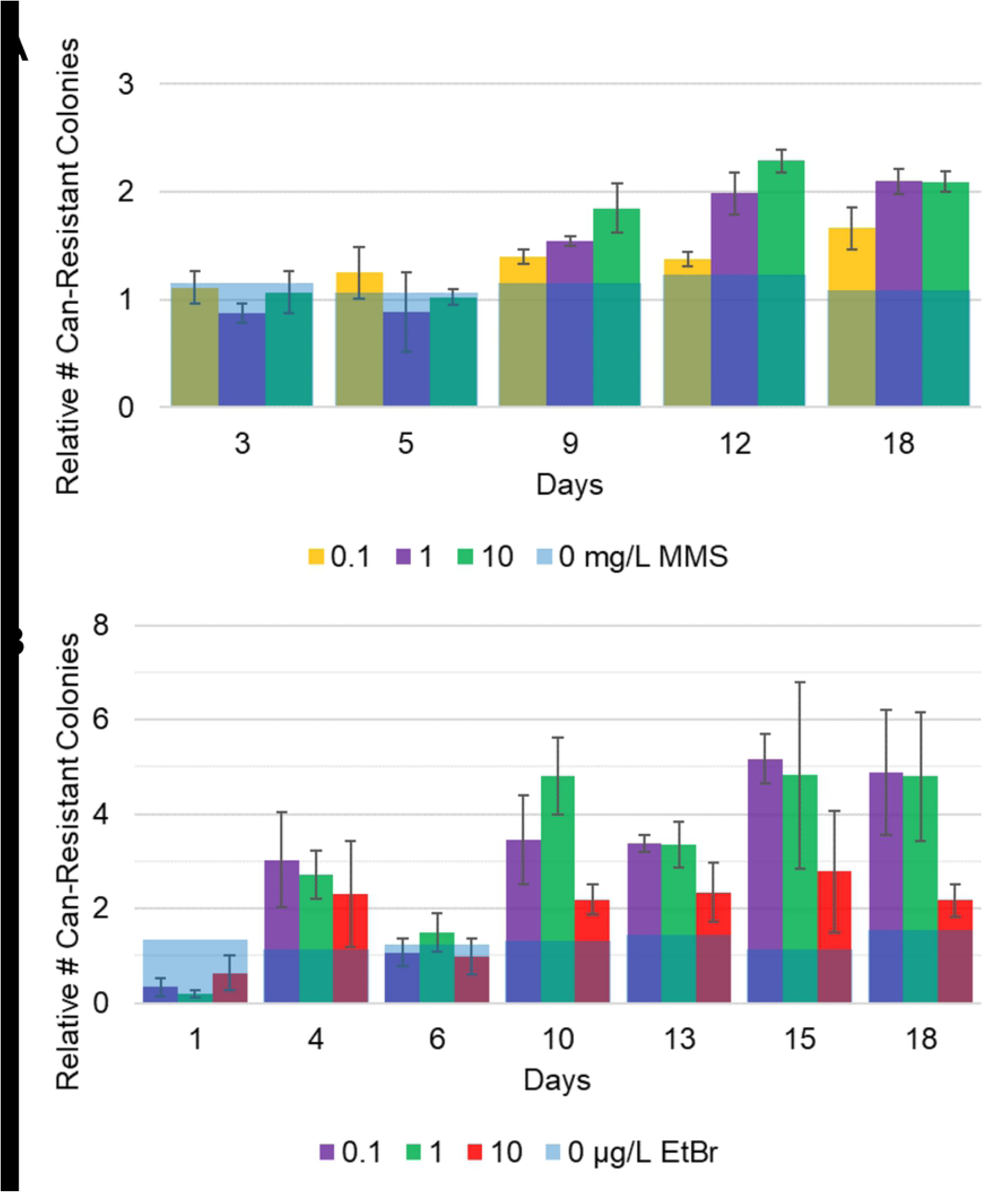
Yeast grown in chemostats can detect levels of mutagen lower than the Ames test via canavanine plate assay. Yeast were grown in parallel chemostats over about three weeks in varying concentrations of the methylating agent MMS (A) or DNA intercalator EtBr (B) and plated on canavanine plates. The number of canavanine-resistant (Can-resistant) colony forming units was counted after 48 hours and graphed as fold-increase relative to the number of Can-resistant colonies from the yeast grown with no mutagen. Each yeast strain was plated in triplicate at each time point. The number of Can-resistant colonies for the no mutagen control is represented as a blue line as the normalized average + normalized standard deviation. For other conditions, the number of Can-resistant colonies is represented as the normalized average +/-normalized standard deviation. The Ames test limit of detections are 2.2 - 4.4 mg MMS/L [32,33] and 2.0 - 31.5 [33,35] μg EtBr/L.

We observed that the number of canavanine-resistant colonies increased with time and with the concentration of mutagen, although not all time points were equally suitable for assessing the level of mutagen. More specifically, it was not possible to distinguish different levels of mutagen until after the first week of growth, after which all concentrations of mutagen tested gave a distinguishable number of canavanine-resistant colonies relative to the untreated condition. During the initial week of culture growth, the number of canavanine-resistant colonies was not clearly distinguishable between each concentration. We speculate this may be due to the time necessary for mutations to accumulate within the sample.

In addition, while the culture continued to grow over the course of the month, the data became noisy after a period of about two and a half weeks, perhaps because of evolution occurring within the population [12,23]. For example, if a cell that is wild-type at the canavanine locus acquires a mutation that enables it to compete significantly better for nutrients in the chemostat environment, then the number of observed canavanine-resistant colonies will drop below that expected for the given level of mutagen. This is a problem that is expected to become more significant with time. Thus, we concluded the data from the yeast chemostats was most accurate from one to three weeks of growth. For this reason, we present the interpretable portion of the data (up to Day 18) in the main text but have provided the full time course in the supporting documents (S**1 Fig**). Moreover, we concluded that we could detect MMS as low as 0.1 mg MMS/mL, about 20 times lower than the Ames test, and that the relationship between the mutagenicity of MMS and its concentration is direct in the range tested, including below the Ames test limit of detection.

### Use of yeast chemostats to measure mutagenicity of the intercalating agent EtBr

To further validate our system, we next grew yeast in the presence of the DNA intercalator ethidium bromide (EtBr), where the Ames test detection limit is 2.0 - 31.5 [33,35] μg/L. We grew yeast for a period of three weeks at 0, 0.1, 1.0, and 10 μg EtBr/L and plated them on canavanine plates as before (**Fig 2B**). Once again, each condition gave a distinguishable number of canavanine-resistant colonies relative to untreated control after about a week. Interestingly, we observed that the lower concentrations of EtBr (0.1 and 1.0 μg EtBr/L) produced higher numbers of canavanine-resistant colonies than the highest concentration of EtBr (10 μg EtBr/L), a biphasic response consistent with hormesis. Briefly, hormesis is when a low dose of a toxin produces a higher effect than a lower dose, perhaps because the higher dose induces protective responses not triggered by the lower dose [36,37]. This observation was reproducible in two more independent runs of the parallel chemostats (S1**C and D Figs**).

We wondered if toxicity of EtBr could account for the observation, even though the concentration of EtBr we used here is far lower than the concentration observed to induce toxicity in *S. typhimurium* (20,000 μg EtBr/L) [21]. To assess the growth of the yeast exposed to EtBr, we tracked culture density and the number of colony forming units on YPD (that is, in the absence of mutagen or canavanine) over time. We saw no significant differences in the culture OD (**S2A Fig**) or in the number of colony forming units per OD unit (**S2B Fig**) in each of the chemostats that would suggest toxicity of EtBr at the concentrations we tested.

In conclusion, similar to MMS, the chemostats could reliably detect the mutagen EtBr as low as 0.1 μg/L, 20 times more sensitive than the Ames test detection limit of 2.0 μg/L (though the chemostats failed to detect the mutagen at 0.01 μg/L, **S2 Fig**). Moreover, unlike the direct relationship seen with MMS, our data provide evidence for hormesis in EtBr. In other words, our data suggest that lower concentrations of EtBr (below those detectable by the Ames test) may have higher mutagenic potential than higher concentrations of EtBr.

### Use of whole-genome sequencing to assess the number of mutations

Next, we asked whether we could use whole genome sequencing as a more sensitive measure of mutation rate. By sequencing and analyzing the whole genomes of individual yeast colonies, we hypothesized that we could get a more sensitive and direct readout of mutagenicity (**Fig 1C**) than from assessing only mutations in a single gene (the *CAN1* locus), as occurs in the plate assay (**Fig 1B**). As shown below, our hypothesis was not correct, at least given the number of colonies we sequenced and the depth of sequence coverage that we used. We present these data and our analysis here so that they may be useful to others attempting to follow up on this work.

From glycerol stocks of yeast grown in the MMS cultures, we isolated the DNA from three independent colonies of yeast grown in chemostats for 9 and 18 days for all four levels of mutagen. We also isolated DNA from three colonies of the yeast used to initially seed the chemostat (“day 0”). We then sequenced the genomes of these yeast for a total of 27 genomes at approximately 10x coverage (see Methods). We originally aligned our reads against the *S. cerevisiae* reference genome for strain S288C (a standard *S. cerevisiae* laboratory strain) or CEN.PK2-1 (the parental strain for our sequenced DBY10148 strain) and used traditional variant calling tools like GATK or VCFtools to identify variants. With this method, however, the number of variants correlated with the number of reads per strain, suggesting that differences arising from inequalities in alignment and coverage were the main contributors to the number of SNPs detected. Moreover, most of the variants detected were differences between our DBY10148 strain and the reference genome strains. To address this issue, we tried generating a *de novo* reference genome using Jellyfish (version 2.2.1) [38] and our Day 0 seed strains, but the coverage of the assembled reference genome was poor, leading to artifacts in data analysis.

To improve our SNP quantification method, we employed somatic mutation calling. Somatic mutation calling, a technique traditionally used to compare sequences from matched tumor and normal tissue samples, compares aligned reads from one sample to another. This technique bypasses inequalities in coverage, as a comparison is only made if both reads have sufficient coverage at a particular locus. Single nucleotide polymorphisms (SNPs) were identified using the somatic mutation caller VarScan (**Fig 3A**) [29]. We used one colony of the initial seed culture as the “normal” sample and one of the remaining 26 genomes as “tumor” samples to identify SNPs caused by MMS exposure and time. As MMS was expected to cause SNPs (and not base pair insertions or deletions [InDels]), we focused our analysis on quantifying the number of SNPs. An analysis of InDels demonstrated no significant difference in the number of InDels between each sample (data not shown).

**Fig 3.**
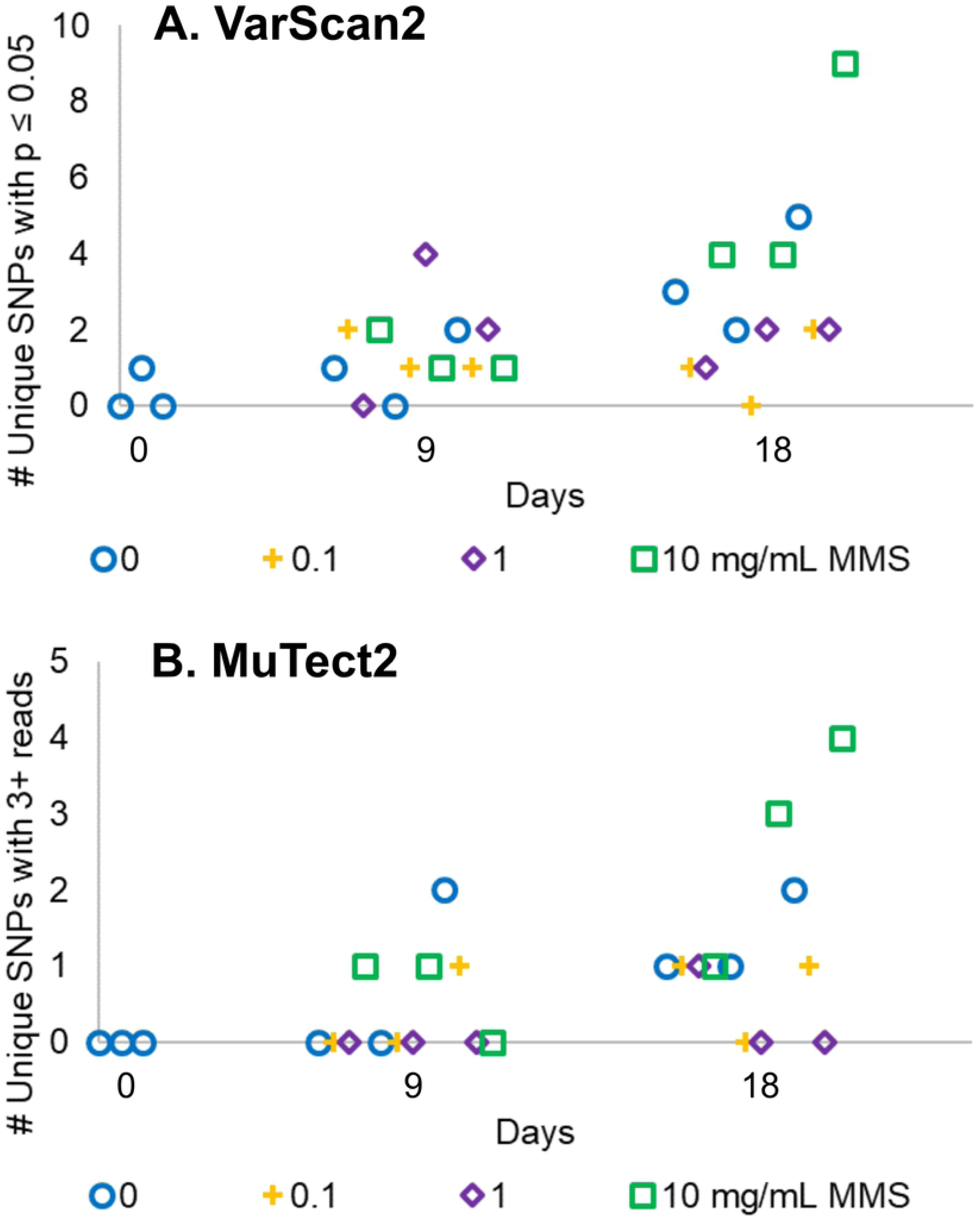
Somatic mutation calling can distinguish yeast grown in high levels of mutagen. Genomic DNA from 27 individual yeast colonies (three per condition) from Fig 2a were sequenced and the number of SNPs quantified, relative to one of the yeast colonies grown in no mutagen at Day 0 using either (A) VarScan2 or (B) Mutect2. Identified SNPs were filtered to remove ambiguous SNPs (SNPs with more than one allele at given genomic position) and a quality check (for VarScan, p value ≤ 0.05 by Fisher’s exact test; for Mutect2, only SNPs with 3 or more reads in “normal” and “tumor” were considered).

Of the approximately 5,000 SNPs identified, the large majority (∼90%) were variant calls with two or more alleles per genomic position, as if the yeast were diploid. This was surprising as the strain DBY1014 is a MATa haploid strain [25]. Calcofluor white staining of log-phase cultures of DBY10148 showed bud scars at only one end (data not shown), further suggesting that the DBY10148 strain indeed is haploid [39]. To understand why sometimes multiple alleles were found at a single genomic position, we analyzed the location and sequence of our diploid-like variants relative to the *S. cerevisiae* S288C reference genome (GenBank GCF_000146045.2; the S288C reference genome is better annotated than the CEN.PK2-1 genome, allowing for easier analysis of SNP location) and saw that these diploid-like variants clustered together (**S3 Fig**). The locations of these diploid-like SNPs corresponded to repetitive DNA sequences like telomeric DNA and viral transposons, similar to what has been previously reported [40]. We reasoned that the alignment method could not accurately map reads corresponding to repetitive DNA sequences [41] and thus disregarded these diploid-like SNPs.

The remaining approximately 500 SNPs that mapped to only one unique genomic location were analyzed to identify variants between the “normal” initial seed culture and the “tumor” mutagenized yeast cultures. Only SNPs that varied between the “tumor” and “normal” with a p-value of 0.05 or lower were counted. Through this process, we identified a number of SNPs that arose due to the spontaneous generation of mutations over time and/or due to the mutagen MMS. However, the number of SNPs identified in yeast cultures grown in 0.1 mg MMS/mL and 1.0 mg MMS/mL were not distinguishable from yeast grown in no MMS (**Fig 3A**). The yeast grown in the highest concentration of MMS, 10 mg/mL, however, was distinguishable from the yeast grown in no mutagen after 18 days in culture.

To confirm these results, we repeated the same analysis with Mutect2, another somatic mutation calling algorithm (**Fig 3B**). Ambiguous SNPs (genome positions with more than one allele) were excluded, and SNPs with two or fewer reads supporting that call were also excluded. Again, we saw that the number of SNPs generally increased over time in continuous culture. Yeast grown in 0, 0.1, and 1.0 mg MMS/mL were not distinguishable at 18 days in culture, but the yeast grown at 10 mg MMS/mL were distinguishable from the no mutagen condition (**Fig 3B**).

## Discussion

Continuous culture systems allow for the growth of a microorganism culture over a long period of time. Here, we demonstrate yeast grown in chemostats can be used as an improved biosensor for mutagenicity. These chemostats were operated for a period of up to a month in the presence of mutagens at different concentrations. The mutagenicity of these environments was then quantified by plating samples of the cultures on canavanine plates, counting the number of colonies that appear, and comparing this number to yeast cultured in media without mutagen. Using this assay, we were able to detect the effects of two established mutagens (the DNA methylating agent MMS and DNA intercalator EtBr) at levels more than one order of magnitude lower than the Ames test. Thus, our yeast assay enables more sensitive detection of mutagenicity of test compounds as well as assessment of the effects of long-term exposure to low levels of established mutagens. Moreover, these devices are relatively cheap and inexpensive to perform and operate, requiring mostly standard laboratory equipment and reagents, such as agar plates.

Variants of the Ames test typically use two additional elements to improve the accuracy of their results. First, the test can be performed with and without the addition of liver extract, as some compounds are not mutagenic but their metabolites or derivatives are [17,42]. Secondly, the typical bacteria used in the Ames test have defects in their DNA repair pathways, allowing for greater sensitivity. While we did not use either of these two elements in our yeast-based system, these may be possible avenues for the improvement of our yeast-based tests for mutagenicity.

The increased sensitivity of our test enabled us to investigate the relationship between the concentration of mutagen and the level of observed mutagenicity at concentrations below the Ames test limit. Typically, this relationship is assumed to be direct, but whether this assumption is accurate has been difficult to assess [43–45]. As the concentration of MMS increased, the number of canavanine-resistant colony forming units increased in a direct manner (**Fig 2A**), consistent with expectation. However, the concentration of EtBr increased, the number of canavanine-resistant colony forming units from yeast grown in 10 μg EtBr/L was fewer relative to yeast grown in lower concentrations (0.1 or 1.0 μg EtBr/L) (**Fig 2B**). This observation was surprising.

To explain the observation that a higher concentration of the EtBr mutagen resulted in fewer mutations, we first hypothesized that the EtBr might have a toxic or growth-inhibiting effect. However, we detected no obvious growth defects in cultures grow in in EtBr as assessed either by optical density of the continuous culture or by the number of colony forming units on YPD (with no mutagen or canavanine) (**S2 Fig**). We cannot rule out that at the highest concentration of EtBr tested may have had an undetected toxic effect on the yeast that would explain our biphasic results. One possible explanation is that higher concentrations of EtBr may have triggered a defense mechanism, such as efflux of EtBr or upregulated DNA repair pathways, that may explain the lower number of canavanine-resistant colonies. Such an explanation would be consistent with hormesis [36,37]. Nonetheless, the number of canavanine-resistant colonies from yeast grown in EtBr from 0.1 to 10 μg EtBr/L was significantly more than control (**Fig 2B**), demonstrating that we could detect the mutagenic potential of yeast even at concentrations lower than the limit of detection for the Ames test. Interestingly, whereas EtBr is only mutagenic in the Ames test in the presence of liver extract [21], we demonstrate that EtBr is mutagenic in yeast without any exogenous metabolic extracts. This aspect of using yeast metabolism may serve as an advantage for using yeast as a biosensor for mutagenicity.

We also tried to assess mutagenic potential via whole genome sequencing. We were originally interested in the sequencing-based approach to quantifying mutagenicity in the continuous cultures because of its potential to be scaled up and automated. While whole genome sequencing was able to detect SNPs in our samples, we observed that the sequencing approach was not as sensitive as the plate-based assay. The data were difficult to interpret for two main reasons. First, for cost reasons, we sequenced at only 10x coverage, and at this coverage and replicate number it was difficult to distinguish SNPs from sequencing errors. Second, the presence of repetitive DNA sequences within the yeast genome made some read alignments ambiguous. Consequently, when a SNP emerged in a repetitive sequence of DNA, it was difficult to distinguish a genuine variant from an artifact of poor alignment. As a result, the overall coverage of our genomic sequences was further reduced because some of the repetitive reads were discarded. In addition, because of cost limitations, we were only able to sequence three individual yeast colonies per condition, which further limited the scope of our analysis.

In retrospect, one might have predicted that the plate-based approach toward determining mutagenicity would be more sensitive because each plate-based approach assays ∼10 million copies of the 1773 base pair *CAN1* gene (the 1 OD unit applied to each plate is approximately 1E7 cells, and many different mutations in the 1773 base pair can result in canavanine resistance), whereas each genome sequence provides information on only the ∼12 million base pair of the yeast genome. Regardless, we are interested in pursuing the sequencing approach because of its potential to overcome the noise (likely due to evolutionary dynamics) observed with the plate assay at long time points (> 3 weeks, **S1 Fig**). A sequencing approach would be strengthened by increasing the read depth, the number of colonies sequenced, and the number of time points sampled, as more data points are needed to obtain the resolution necessary to distinguish between the low and medium concentrations of MMS. However, sequencing costs would increase. Moreover, improved methods of mapping repetitive regions may improve the sensitivity assessing mutagenicity by sequencing. Thus, practical application of the sequencing approach may await further improvements in sequencing technology and bioinformatics. In the meantime, the plate assay remains straightforward and practical for exposure times up to ∼3 weeks and provides a ∼20x improvement over the sensitivity of the Ames test.

In conclusion, we present a method for the long-term continuous culture of yeast as an improved biosensor for mutagenicity. We have shown that this biosensor works for mutagens that act by different mechanisms and suggest that it might have particular utility for assessing the safety of compounds (e.g. food additives, trace contaminants in reclaimed water) to which humans are exposed at low concentrations for long periods of time.

## Acknowledgements

We thank the staff of the Notre Dame Genomics & Bioinformatics Core Facility, particularly Melissa Stephens, the Notre Dame Center for Research Computing, and Prof. Scott Emrich for their support. We also thank many bioinformaticians for their advice and troubleshooting. This work was partially supported by NSF DBI 1556349 to HVG. The sequencing and bioinformatics efforts were supported by a grant to HVG from the Indiana CTSI. JYO received support from the Glynn Family Honors Program. JTP received support from the University of Notre Dame Chemistry, Biochemistry, Biology Interface Program (NIH T32GM075762), American Water Works Association Thomas R Camp Fellowship, and the University of Notre Dame Center for Aquatic Conservation Fellowship. The funders had no role in study design, data collection and analysis, decision to publish, or preparation of the manuscript.

S1_Table.xls

Contains the source data (individual numbers) for data in Fig 2, Fig 3, SFig 1, and SFig 2.

**S1 Fig. Complete and additional timecourses of MMS and EtBr chemostats**.

(A) Complete timecourse of MMS chemostats shown in Fig 2A.

(B) Complete timecourse of EtBr chemostats shown in Fig 2B.

(C, D) Two additional replicates of EtBr chemostats performed in a similar manner to Fig 2B. While yeast grown in chemostats with 0.1, 1.0, and 10 µg EtBr/L consistently produced more canavanine-resistant colonies compared to the no mutagen control, the concentration 0.01 µg EtBr/L was unable to be reliably detected. Moreover, the concentration 1.0 µg EtBr/L consistently produced more canavanine-resistant colonies than 10 µg EtBr/L. Data are presented as in Fig 2.

**S2 Fig. Low levels of EtBr do not cause toxicity to yeast grown in continuous culture systems**. To determine if EtBr had significant toxicity to the yeast, (A) the OD600 of the yeast was assessed over time to identify growth defects and was determined to have no significant variation. (B) To determine if there was any difference in the number of colony forming units, 1.25E-6 OD units of yeast (roughly 40 cells) were obtained by serial dilution and plated onto YPD plates in triplicate. The number of colony forming units (CFU) were counted after 48 hours at 30°C and presented as the average and standard deviation. The data here correspond to the EtBr chemostats presented in Sup. Fig. 2D.

**S3 Fig. Diploid-like SNPs cluster in genomic regions**. The location of SNPs (colored circles) found via VarScan by comparing reads from a non-mutagenized colony of DBY10148 to the S288C reference genome were visualized according to their genomic position. Yeast chromosomes 1 to 16 are represented by a solid black line and are numbered. The mitochondrial genome is denoted as chromosome 17. A red dot approximates the centromeric region.

